# Single-molecule imaging reveals the interplay between transcription factors, nucleosomes, and transcriptional bursting

**DOI:** 10.1101/404681

**Authors:** Benjamin T. Donovan, Anh Huynh, David A. Ball, Michael G. Poirier, Daniel R. Larson, Matthew L. Ferguson, Tineke L. Lenstra

## Abstract

Transcription factors show rapid and reversible binding to chromatin in living cells, and transcription occurs in sporadic bursts, but how these phenomena are related is unknown. Using a combination of *in vitro* and *in vivo* single-molecule imaging approaches, we directly correlated binding of the transcription factor Gal4 with the transcriptional bursting kinetics of the Gal4 target genes *GAL3* and *GAL10* in living yeast cells. We find that Gal4 dwell times sets the transcriptional burst size. Gal4 dwell time depends on the affinity of the binding site and is reduced by orders of magnitude by nucleosomes. Using a novel imaging platform, we simultaneously tracked transcription factor binding and transcription at one locus, revealing the timing and correlation between Gal4 binding and transcription. Collectively, our data support a model where multiple polymerases initiate during a burst as long as the transcription factor is bound to DNA, and a burst terminates upon transcription factor dissociation.

## Introduction

During activation of transcription, transcriptional factors (TFs) bind to specific motif sequences in the promoters of genes and recruit factors such as chromatin regulators, coactivators, general transcription factors and eventually RNA polymerase to initiate transcription. Understanding the molecular events that underlie gene activation requires knowledge about the kinetics of these processes. Studies of transcription dynamics in single cells have shown that some genes are constitutively transcribed with random initiation of single polymerases (1-state model) (Larson et al., 2011; Zenklusen et al., 2008), whereas other genes are transcribed in short stochastic bursts of high transcriptional activity where several polymerases initiate, interspersed by periods of inactivity (2-state model) (Chubb et al., 2006; Golding et al., 2005; Lenstra et al., 2015; Bahar Halpern et al., 2015; Zenklusen et al., 2008). Modulation of a gene’s transcriptional activity can be done by changing the burst size (the number of polymerases in a burst), the burst duration or the burst frequency (Bartman et al., 2016; Dar et al., 2012; Fukaya et al., 2016; Molina et al., 2013), each with different effects on the cell-to-cell variability in a population (Raser and O’Shea, 2004).

Recent advances in imaging technologies allow for the direct measurement of TF binding dynamics at the single molecule level in living cells, providing insight into the search dynamics and mechanisms of gene activation (Elf et al., 2007; Tokunaga et al., 2008; Grimm et al., 2015; Liu and Tjian, 2018). Interpretation of the TF dynamics has been challenging, because the position and activity of target genes are often unknown, and target genes have different TF binding site affinities (Normanno et al., 2015). Target genes are also in different chromatin contexts, where the position or modification state of promoter nucleosomes can affect the accessibility of binding sites for TFs, although certain TFs, such as pioneer TFs, bind to DNA even in the context of a full or partially unwound nucleosome (Zaret and Mango, 2016). For example, at the *GAL1*/*10* locus in budding yeast, the TF Gal4 binds to a partially unwrapped nucleosome with the help of the RSC remodeler (Floer et al., 2010). In addition to regulating accessibility, nucleosomes can significantly reduce the dwell time of transcriptional activators *in vitro* (Luo et al., 2014). However, how nucleosomes regulate TF binding dynamics *in vivo* is still mostly unexplored. In addition, the relationship between TF binding dynamics and the dynamics of transcription initiation is only starting to emerge (Larson et al., 2013; Senecal et al., 2014; Rullan et al., 2018; Loffreda et al., 2017).

The major difficulty in deciphering the causal relationships between TF dwell time, nucleosomes and the dynamics of RNA synthesis comes from technical limitations. Although TF kinetics and RNA synthesis can be measured at single-molecule resolution, it has been challenging to measure both simultaneously in the same cell at the active locus. Even if the location of target genes is known, the number of TF binding events at a specific locus would be limited, because of the high number of target genes and partial labelling of the TF population in single-molecule imaging experiments (Liu and Tjian, 2018). Additionally, there is a mismatch in the time scale of TF binding (on the order of seconds) and transcription (on the order of minutes), and quantification of the transcription site intensity in a single plane is hampered by transcription site movement. Similarly, assessing the role of nucleosomes in regulating TF and transcription dynamics has proven difficult, since we lack *in vivo* tools to precisely control or visualize the position of nucleosomes around TF binding sites in single cells and observe their effect on TF binding dynamics.

Here, we used a combination of novel single-molecule *in vitro* and *in vivo* imaging techniques to assay the interplay between TF dwell time, nucleosome binding and transcriptional bursting. We have developed a novel single-molecule imaging platform to directly visualize both transcription binding dynamics and transcriptional output in the same cell in an endogenous setting. We exploited the characteristics of the galactose responsive genes in budding yeast, where the combination of low Gal4 expression and a small amount of Gal4 target genes allows for the quantification of binding events to the specific target gene of interest. Moreover, we have employed an advanced 3D tracking technique to track the transcription site in 3D, which has allowed for the first time to directly correlate TF binding with transcription initiation kinetics at a single locus. We find that Gal4 precedes and overlaps with *GAL10* transcription, and that their fluctuations are coupled in time. Gal4 dwell time determines the transcription burst duration and depends on the affinity of the binding site. The burst duration is not modulated by galactose signaling, which instead regulates burst frequency. In addition, quantitative comparison of the *in vivo* Gal4 dwell time to the *in vitro* Gal4 binding rates in a nucleosomal context indicate that promoter nucleosomes reduce the Gal4 dwell time by orders of magnitude allowing for rapid Gal4 turnover. Overall, we show that TF binding to nucleosomal DNA is a key determinant of transcriptional burst duration, where multiple transcription initiation events can take place as long as the TF is associated with DNA.

## Results

### Mutations in the upstream activating sequences reduces burst size

To study the role of TF binding in bursting, we focused on the *GAL3* gene, which contains one Gal4 UAS (upstream activating sequence) in its promoter. Endogenous *GAL3* transcription was visualized in live cells by the introduction of 14 PP7 repeats in the 5’ UTR of *GAL3*. Upon transcription, each PP7 repeat forms a stem loop that is bound by the PP7 bacteriophage coat proteins fused to a fluorescent protein. The PP7 loops did not affect *GAL3* transcription levels, as the transcription site (TS) of the PP7-tagged and non-tagged allele in heterozygous diploid cells showed similar amount of nascent RNAs (Figure S1A).

Live-cell visualization of *GAL3* transcription in galactose-containing media revealed transcriptional bursts, with periods of high activity interspersed with periods of inactivity (Figure 1A, movie S1). The bursts were less frequent than for the previously measured *GAL10* gene (Lenstra et al., 2015), with longer periods of gene inactivity. To determine on and off periods of transcription, a threshold was applied to intensity traces of the transcription site. In agreement with a two state-model of bursting (Peccoud and Ycart, 1995), both the on and the off time are exponentially distributed (Figure 1B and C), with an average on time of 46.5s ± 2.4s and average off time of 4.2 ± 0.2 min.

**Figure 1.**
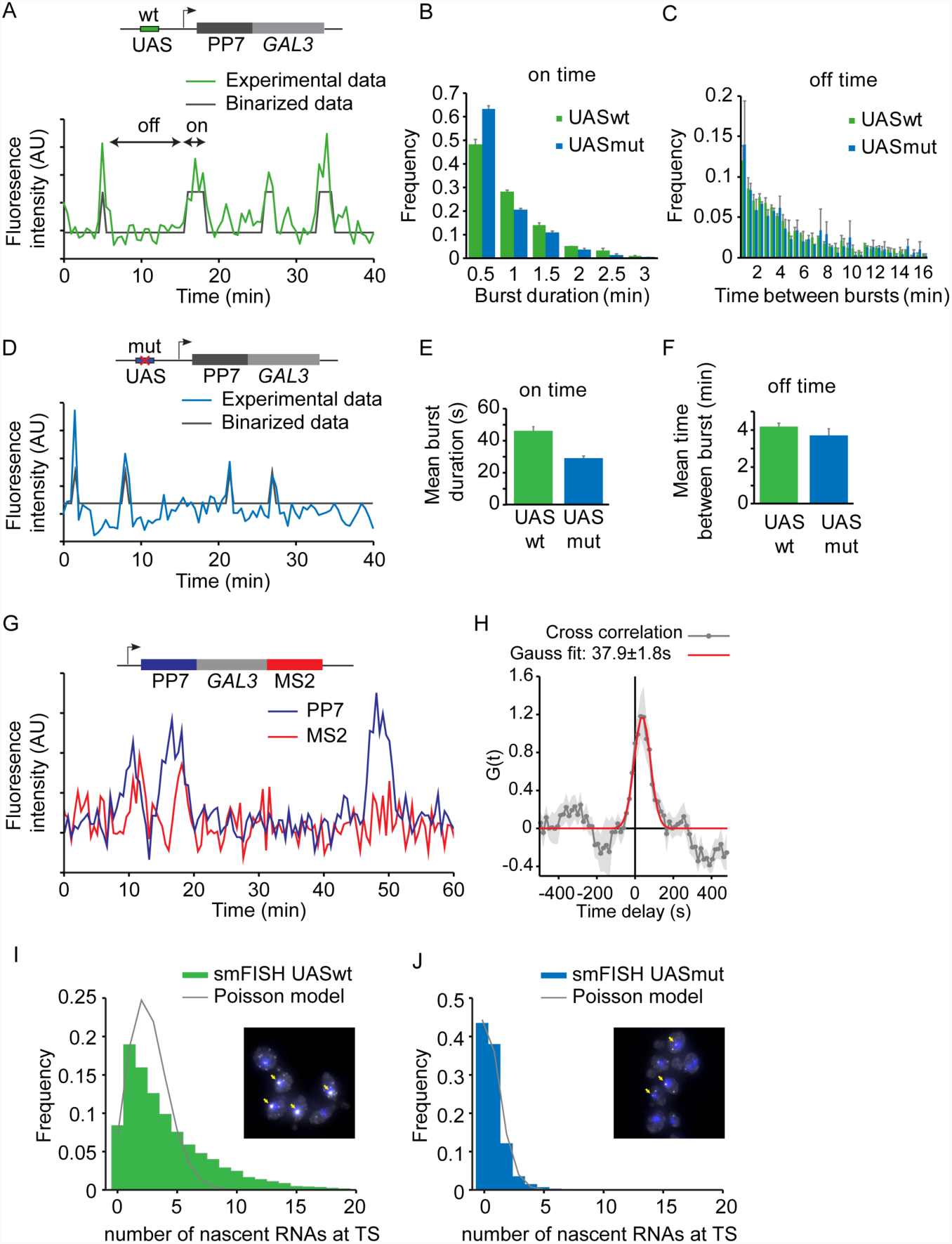
Mutations in upstream activating sequence (UAS) reduces burst size, but not burst frequency. (A) Transcription at *GAL3* is visualized by addition of PP7 loops at the 5’ of *GAL3*. Example trace of the quantified fluorescence intensity of the transcription site. Traces are binarized to determine on (active) and off (inactive) times. (B) Histogram of *GAL3* on time (burst duration) for cells with wt and mutated UAS, respectively, reveals shorter on times for mutated UAS. Errors indicate SE of 3 experiments. (C) Histogram of *GAL3* off times. Errors indicate SE of 3 experiments. (D) Similar to (A), but from cells were the UAS was mutated to a lower affinity UAS. Also see Figure S1. (E). Average on time for UASwt and UASmut from exponential fit. Errors indicate SD of fit. (F) Average off time for UASwt and UASmut from exponential fit. Errors indicate SD of fit. (G) Example traces of quantified fluorescence intensity at the transcription site for 5’PP7 and 3’MS2 at *GAL3.*(H) Cross correlation of the PP7 and MS2 signals shows a peak at 37.9s±1.8s delay (I) Distribution of number of nascent RNAs at the transcription site determined by smFISH. The distribution of UASwt does not fit a Poisson distribution (grey line), supporting that *GAL3* is not constitutively transcribed, but is transcribed in bursts. Example image in shown, arrows indicate TSs. (J). Same as (G) for cells with UASmut. The distribution of UASmut fits a Poisson distribution, indicating that GAL3-UASmut is transcribed with random initiation of individual polymerases, similar to constitutive genes.

To investigate the role of Gal4 in determining this bursting pattern, the *GAL3* UAS upstream of the PP7-*GAL3* gene was replaced with one of the *GAL10* UAS sequences with a lower affinity (called UASmut, Figure 1D) (Liang et al., 1996). Interestingly, analysis of the traces in this mutant revealed that the on time was significantly reduced from 46.5s ± 2.4s to 29.3s ± 1.3s, whereas the off times were similar (4.2 ± 0.2 min for UASwt and 3.7 ± 0.3 min for UASmut, Figures 1 C-F). The on time of transcription (also referred to as burst duration) reflects the time that nascent RNA is visible at the site of transcription and includes the window of time when polymerases are loaded on the gene as well as the duration of elongation, termination, and release. We refer to the ‘active promoter time’ as the interval over which RNA polymerases initiate at the promoter and the ‘burst size’ as the number of polymerases which are loaded during this period. If post-initiation processes are similar for both UASwt and UASmut, a reduced on time indicates that the active promoter time is reduced, likely resulting in fewer polymerases initiating during a burst for the UASmut.

To distinguish these different steps in the transcription cycle, we inserted the PP7 loops in the 5’ UTR of *GAL3* and the orthogonal MS2 loops in the 3’ UTR of *GAL3*. As expected, dual color imaging of this PP7-*GAL3*-MS2 strain showed an increase in PP7 intensity followed by an increase in MS2, and a simultaneous drop of both signals (Figure 1G, movie S2). The peak in the PP7-MS2 temporal cross correlation revealed that RNA polymerase takes on average 37.9s ± 1.8s to transcribe from the middle of the PP7 to the middle of the MS2 loops (Figure 1H), with an elongation rate of 64.5 ± 3.0 bp/s (3.9 ± 0.2 kb/min). Based on the length of the PP7-*GAL3* construct, the elongation time of this construct is calculated to be ~30s. The average on time of 29.3s ± 1.3s in the UASmut construct thus suggests that the transcription on time is dominated by the elongation time with little time for loading of multiple polymerases. Therefore, the UASmut facilitates transcription of single RNAs during a burst with an active promoter time of a few seconds, whereas UASwt has a longer active promoter time of ~ 15 s.

To confirm that the burst size is reduced upon mutating the UAS, smFISH was performed on the UASwt and UASmut strains (Figure 1I and J). The distribution of nascent *GAL3* RNA normalized to intensity of cytoplasmic RNAs, shows that multiple RNAs are present at the TS in a *GAL3* UASwt strain. In agreement with bursting transcription, the nascent transcript distribution does not fit to a Poisson model, which is the expected distribution for non-bursting constitutive genes. In the strain with a UAS mutation, the number of nascent transcripts is reduced and fits well to a Poisson model. The UAS mutation thus results in a reduction in the burst size to approximately one RNA per burst, similar to a constitutively transcribed gene. In summary, bursts of *GAL3* transcription require high affinity binding of Gal4 to the promoter and reducing the affinity of Gal4 through a cis-acting mutation in the UAS reduces the burst size.

### Mutation in the UAS reduces dwell time of Gal4 on nucleosomal DNA *in vitro*

Since the *in vivo* experiments suggest the decrease in burst duration is due to a decrease in the affinity of Gal4 for the UAS, we performed a series of *in vitro* binding assays to directly compare Gal4 binding to the UASwt and UASmut sequences. First, we compared Gal4 binding affinity to the two sequences on naked DNA. In these experiments, Gal4 occupancy on naked DNA was measured with protein induced fluorescent enhancement (PIFE)(Hwang et al., 2011), where a Gal4 binding event enhances the fluorescence of a Cy3 fluorophore ~1.5 fold (Figure 2A). Titrating Gal4 produces a binding curve that indicates stoichiometric binding to the UASwt sequence (Figure S2A). Because of the stoichiometric binding, the relative affinity of Gal4 to the UASwt and UASmut sequences could be determined by titrating different concentrations of unlabeled competitor DNA. Wildtype competitor DNA was more effective in removing Gal4 from the labeled DNA than mutant competitor (IC_50_ 4.0 ± 0.6 nM for UASwt vs 17.3 ± 3.5 nM for UASmut, Figure 2B), with a relative affinity change of 4.3 fold (Figure 2C).

*In vivo*, DNA is wrapped around nucleosomes. Previous measurements of Gal4 binding on naked versus nucleosomal DNA showed that nucleosomes reduce the dwell time of Gal4 ~1000 fold (Luo et al., 2014). Binding of Gal4 to the UAS sequences was therefore measured within a nucleosome by determining the change in energy transfer between two fluorophores positioned in the nucleosome entry/exit and DNA (Förster resonance energy transfer, FRET). Gal4 binding traps the nucleosome in a partially unwrapped state, which increases the distance between the fluorophores, reducing the FRET efficiency (Figure 2D). Similar to naked DNA, Gal4 has an 8.6 fold higher affinity for the UASwt relative to the UASmut sequence (Figure 2C and E).

**Figure 2.**
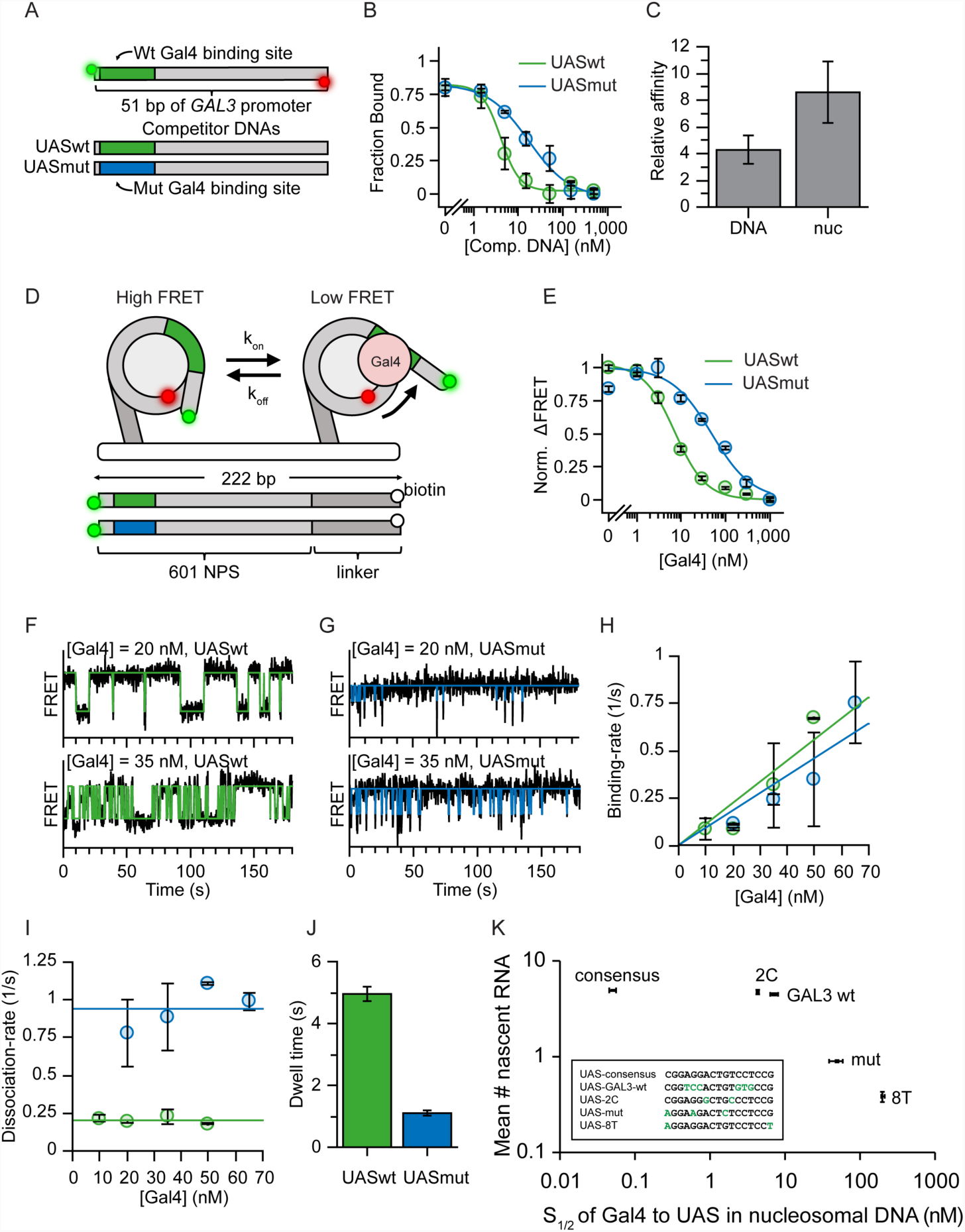
Mutations in UAS reduce residence time of Gal4 on nucleosomal DNA *in vitro*. (A) Competitive binding experiments were performed to determine relative affinity of Gal4 to UASwt (green) and UASmut (blue) motifs. Gal4 occupancy on UASwt Cy3/Cy5 DNA was determined by measuring protein induced fluorescence enhancement (PIFE) on 51 bp oligos containing either Gal4 UASwt or UASmut site 1 bp away from Cy3 fluorophore). (B) Titrating unlabeled UASwt or UASmut competitor DNA reduces UASwt Cy3/Cy5 DNA occupancy. IC_50_UASwt = 4.0 ± 0.6 nM, IC_50_UASmut = 17.3 ± 3.5 nM. (C) Comparison between relative affinities of Gal4 UASwt versus UASmut in naked or nucleosomal DNA shows 4.3 ± 1.1x difference in KD from naked DNA and 6.8 ± 1.7x from nucleosomal DNA. (D) Experimental setup for smFRET experiments to measure Gal4 binding at nucleosomal DNA. Gal4 binds to site 8 bp into nucleosome. A FRET pair in the entry/exit region provides readout on binding events. In the absence of Gal4, nucleosomes are in the high FRET state. A Gal4 binding event traps nucleosome in low FRET state. (E) Gal4 affinity to nucleosomes containing UASwt or UASmut sequences gives S_1/2_ of 7.2 ± 0.8 nM and 48.9 ± 10.8 nM, respectively. (F) Example smFRET traces showing Gal4 binding to UASwt in nucleosomal DNA at two different Gal4 concentrations. States are determined using HMM fit. (G) Same as G but for UASmut. (H) Binding-rates of Gal4 are concentration dependent, but are similar for UASwt and UASmut: k_on_ UASwt = 0.011 ± s^-1^ nM^-1^, k_on_ UASmut = 0.009 ± 0.001 s^-1^ nM^-1^ (I) The dissociation-rate of Gal4 at UASwt is ~5-fold slower compared to the UASmut: k_off_ UASwt = 0.20 ± 0.01 s^-1^, k_off_ UASmut = 0.92 ± 0.05 s^-1^. (J) Histogram of Gal4 dwell time to nucleosomal DNA containing UASwt and UASmut sequences. (K) Scatter plot showing the average number of nascent RNA at TS (from smFISH) vs affinity of Gal4 to different UAS sequences in nucleosomal DNA (from in vitro measurements). *In vivo* transcription levels correlate with *in vitro* affinity of Gal4, but transcription saturates above wildtype sequence. UAS sequences are shown in the box.

We then directly measured the binding and dissociation rates of Gal4 to both UAS sequences in a nucleosomal context by recording time-resolved single-molecule FRET trajectories between Gal4 and surface-tethered nucleosomes (Figure 2F and G). As expected, the binding-rate (frequency of binding) but not the dissociation-rate (dwell time) is Gal4 concentration dependent (Figure 2H and I). The binding rates are similar for UASwt and UASmut, indicating that the frequency of binding is independent of the UAS sequence and likely regulated by the nucleosome unwrapping rate (Figure 2H). Interestingly, the UASmut shows a 5-fold shorter dwell time than the wildtype (4.96s ± 0.25 for UASwt, 1.09s ± 0.06 for UASmut, Figure 2I and J). A higher affinity binding site thus stabilizes Gal4 binding but does not influence the frequency of binding. This higher dwell time correlates with an increased burst size *in vivo*.

To generalize the relationship between binding affinity and transcription, we measured the affinity of five different sequences to nucleosomal DNA *in vitro* in comparison to the mean number of nascent RNAs *in vivo*, as determined by smFISH (Figure 2K). In agreement with our previous data, lower affinity binding correlates with lower nascent RNA production. However, binding sites with higher affinity than the wildtype sequence did not increase nascent RNA output. This saturation suggests that either other regulatory factors limit the Gal4 dwell time *in vivo*, or there is an inherent limit to the burst size even upon longer Gal4 binding.

To test whether *in vitro* measurements of Gal4 binding to nucleosomal DNA have *in vivo* relevance, we sought to determine the nucleosome occupancy near Gal4 binding sites in the presence or absence of galactose, i.e. induced and uninduced conditions. Specifically, we mapped stable and fragile nucleosomes by digestion with two different MNase concentrations (Kubik et al., 2015). Cells grown in galactose showed protected fragments of nucleosomal size (95-225 bp) around the Gal4 binding site in the *GAL3* promoter (Figure 3). This MNase peak was only visible with low MNase digestion and only appeared after the two flanking stable nucleosomes moved up- and downstream, which suggests the presence of a fragile nucleosome around the Gal4 UAS (Kubik et al., 2015). Fragile nucleosomes have also been reported around the *GAL1-10* UAS sites (Floer et al., 2010), and we find that fragile nucleosomes appear at many other galactose responsive genes (Figure S3), strongly suggesting that Gal4 binds nucleosomal instead of naked DNA *in vivo*.

**Figure 3.**
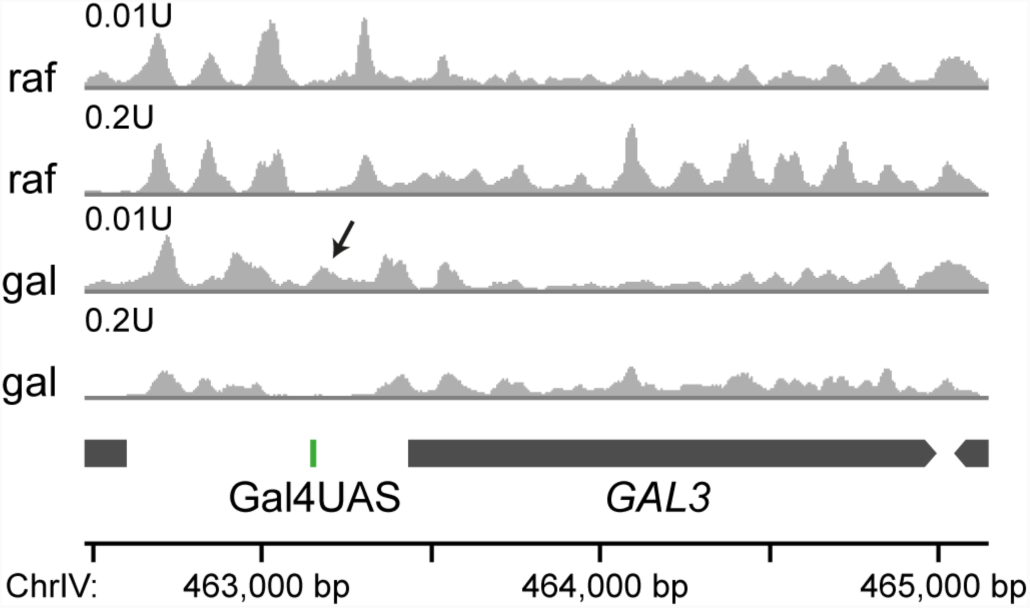
Gal4 binds at the edge of a fragile nucleosome in galactose *in vivo*. Profiles of nucleosome midpoint positions by MNase-seq experiments. Samples were digested with the indicated MNase concentrations in both raffinose (raf) and galactose (gal) containing media. Midpoints of nucleosomes are smoothed by 31 bp. In galactose the stable nucleosomes move away from the Gal4UAS, creating space for an additional fragile nucleosome (indicated by arrow).

Overall, *in vitro* and *in vivo* data indicate that Gal4 binding dynamics are correlated with RNA production, that longer binding supports larger burst sizes, that dwell time is shorter on nucleosome-bound DNA, and that Gal4 binding sites contain fragile nucleosomes.

### Gal4 dwell time *in vivo* shows two Gal4 binding populations

Our *in vitro* dwell time measurements indicate that Gal4 is bound to nucleosomal DNA on the order of seconds. Previous Gal4 binding measurements to naked DNA showed a 1000 fold longer dwell time (Luo et al., 2014), suggesting that nucleosomes may be required for rapid turnover of TFs in the nucleus. In living cells, many additional factors may also limit the dwell time of Gal4. We therefore used single-molecule tracking (SMT) to analyze the Gal4 dwell time *in vivo*.

To visualize binding and diffusion of single molecules of Gal4, Gal4 was tagged with a HALO tag, which covalently binds bright synthetic dyes (Figure 4A). Addition of the HALO tag minimally affected its function, as Gal4-HALO cells showed similar growth as wildtype on galactose-containing plates (Figure S4A). To support uptake of the synthetic dye, the gene encoding the Pdr5 transporter was also deleted (Ball et al., 2016). Lastly, 14 PP7 loops were added in the 5’UTR of *GAL10* (Figure 4A), allowing the visualization of Gal4 binding kinetics specifically at one of its target genes. For this experiment we used *GAL10* instead of *GAL3* because we reasoned that higher transcriptional activity may increase the chance of observing colocalization of TF binding at its target gene with respect to nascent RNA production.

**Figure 4.**
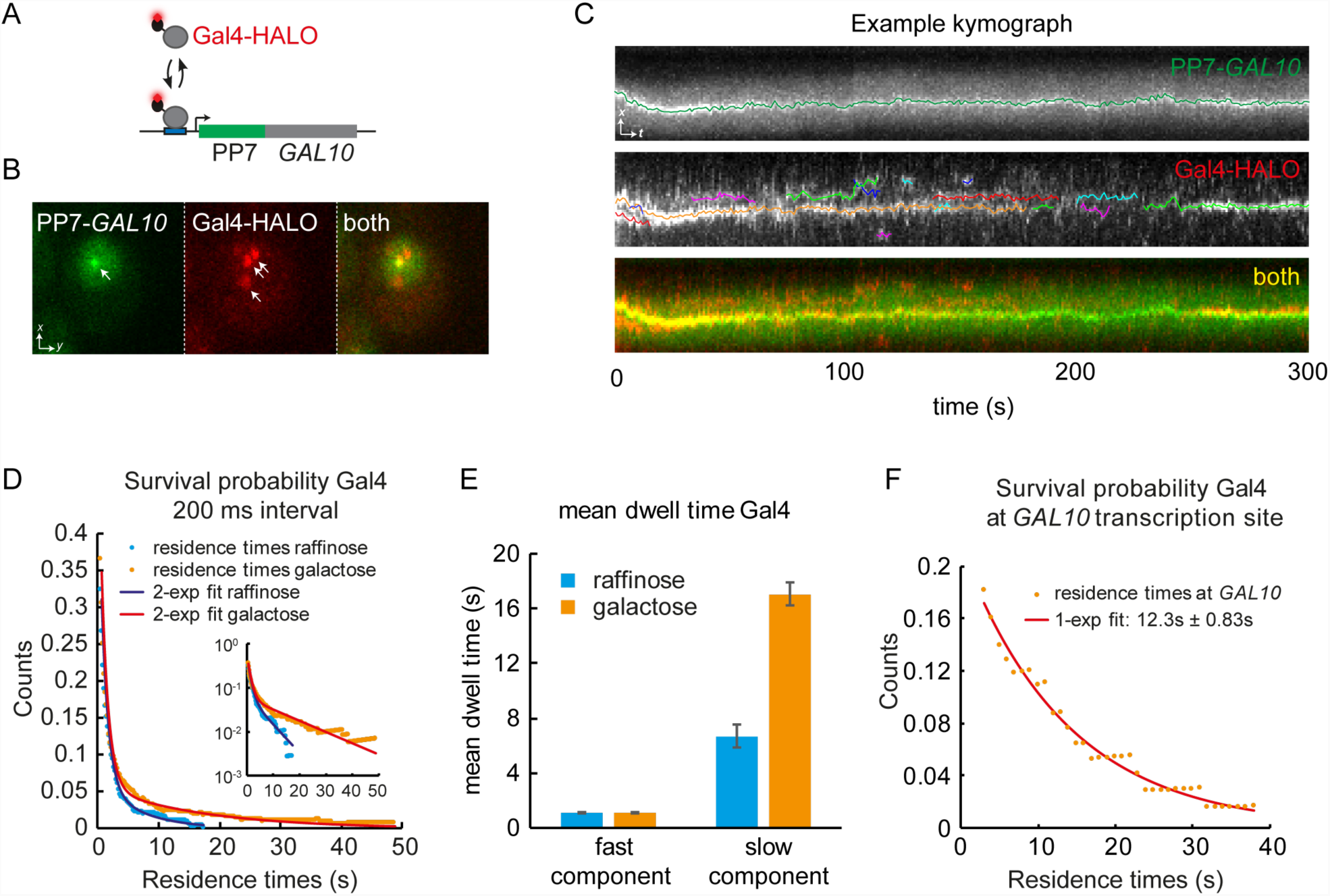
Measurements of the Gal4 residence time *in vivo* shows two Gal4 populations, with long residence times colocalizing with target gene. (A) Simultaneous imaging of Gal4 binding kinetics *in vivo* using single-molecule tracking and RNA imaging of the *GAL10* target gene. Gal4 is tagged with a HALO-tag, which covalently binds to the dye JF646. Transcription of the target gene *GAL10* is visualized by PP7-loops. (B) Example image of a cell, showing the *GAL10* TS (arrow, left panel), single molecules of Gal4 (arrows, middle panel) and an overlay (right panel). (C) Example kymograph of a cell. Upper panel (PP7-*GAL10*) shows the position of the TS over time. Middle panel (Gal4-HALO) show tracks of Gal4. Data is in white, colored lines show analyzed tracks. Lower panel shows overlay, showing colocalization of some the Gal4 tracks to the *GAL10* TS. (D) Survival probability of the duration of Gal4 tracks (after displacement thresholding) at 200 ms interval from cells grown in raffinose or galactose. Lines show bi-exponential fit, indicating 2 Gal4 populations. Inset shows data in semi-logarithmic plot. (E) Residence time of the fast and slow component of the fits from (D). The slow component changes between conditions. (F) Survival probability of Gal4 residence times that colocalize with *GAL10* TS (<250nm), showing that Gal4 with long residence time colocalizes with *GAL10*. Data was taken at 1s interval. Line shows exponential fit, with mean of 12.3s ± 0.83s.

Cells were imaged with two colors simultaneously using HILO illumination (Highly Inclined and Laminated Optical sheet) to reduce out-of-focus fluorescence (Tokunaga et al., 2008). We observed single Gal4 molecules in the nucleus, some of which colocalize with the PP7-*GAL10* TS (Figure 4B, movie S3). The *GAL10* TS and Gal4 molecules were tracked over time, as shown in the kymograph of Figure 4C. To determine if Gal4 molecules were diffusing or bound to DNA, a distance threshold between frames was applied based on the mobility of histone H3 (*HHT1*-HALO) molecules.

The survival probability of bound Gal4 molecules at all sites reveals binding of Gal4 on the order of several seconds, which is in the same dynamic range as the *in vitro* measurements on nucleosomal DNA (Figure 2J and (Luo et al., 2014)). The biphasic behavior in the semi-log plot indicates two populations of Gal4 with different binding rates (inset, Figure 4D). Multiple distributions have previously been observed for other TFs and have been attributed to non-specific and specific binding (Ball et al., 2016; Chen et al., 2014; Presman et al., 2017). In agreement with this interpretation, the core histone H3 only showed a longer binding population and no shorter binding population (Figure S4B). A bi-exponential fit of the Gal4 survival probability revealed that the fast binding fraction of ~1s has a similar dwell time in inactive and active conditions (raffinose and galactose, respectively, Figure 4E), supporting the idea that the fast binding is non-specific. In contrast, the dwell time of the more stable binding fraction increases from 6.7s ± 0.9s to 17.1s ± 0.8s in active conditions. Since galactose responsive genes are inactive or lower transcribed in raffinose than in galactose, an increased dwell time suggests that longer binding is specific binding that is required to activate transcription.

Interpretation of TF dwell times in the complex milieu of the nucleus can be challenging since residence times are averaged across many binding sites that likely have different affinities. To obtain insight into the binding kinetics at a single target gene, we took advantage of the addition of PP7 RNA labeling for the *GAL10* gene. When we focused on Gal4 binding events that specifically overlapped with the *GAL10* transcription site in the presence of galactose, several longer binding events were observed, with a mean dwell time of 12.3s ± 0.83s (Figure 4F). Since alignment between channels may be imperfect, our stringent colocalization threshold of 250nm may result in underestimation of the dwell time at *GAL10*. Regardless, the presence of longer binding events at a transcribing target gene indicates that more stable binding is associated with the activation of transcription.

The dwell time of TFs can be underestimated if molecules are bleached before they dissociate from the DNA. Histone H3 binds on much longer time scales and served as a control to determine the maximally measurable dwell time. Since experiments taken with 200 ms interval showed Gal4 dwell times similar to H3 dwell time, the same experiments were repeated with 1s interval. At this longer time interval, non-specific binding was below the time-resolution, and the survival probably followed a single exponential curve. Although the longer interval indeed resulted in an increase of H3 dwell time, Gal4 dwell time did not increase significantly (Figures S4B-D). The measured average specific Gal4 dwell time (~17 s) is therefore unlikely to be the result of a technical artifact. Moreover, the fraction of Gal4 associated with DNA also increases in active versus inactive conditions (Figure S4E), in agreement with more binding sites becoming available.

In summary, Gal4 shows two binding populations. Our data supports a model where the short sub-second binding is non-specific binding of Gal4 in search for its target sites, and that the more stable several-second binding is specific and productive binding to the promoters of target genes in order to activate transcription.

### Gal4 binding correlates with transcription

Long Gal4 binding events (~ 12 – 17 s) which spatially co-localize with the *GAL10* TS suggests that these binding events result in the production of nascent RNA. However, the SMT experiments visualizes molecules in a single plane, and the three-dimensional movement of the *GAL10* TS out of the focal plane hampered quantification of the TS intensity. Using SMT, we were therefore unable to quantify if binding was followed by the production of RNA. Moreover, there is a dynamic mismatch which poses a challenge for fluorescence visualization: Gal4 is bound on the order of 10 seconds, but bursts of transcription are separated by many minutes. To directly correlate Gal4 binding with RNA output, we imaged our dual tagged Gal4-HALO and PP7-*GAL10* cells in a 3D orbital tracking microscope (Levi et al., 2005a, 2005b), which is able to track the *GAL10* gene in 3D and simultaneously capture Gal4 binding. As a variation on conventional confocal scanning microscopy, an orbital tracking microscope makes circular orbits around an object of interest and uses a feedback loop to track the object in 3D with high temporal and spatial resolution (Figure 5A). Figure 5B shows an example trace of peaks of Gal4 binding, followed by increases in *GAL10* transcription.

**Figure 5.**
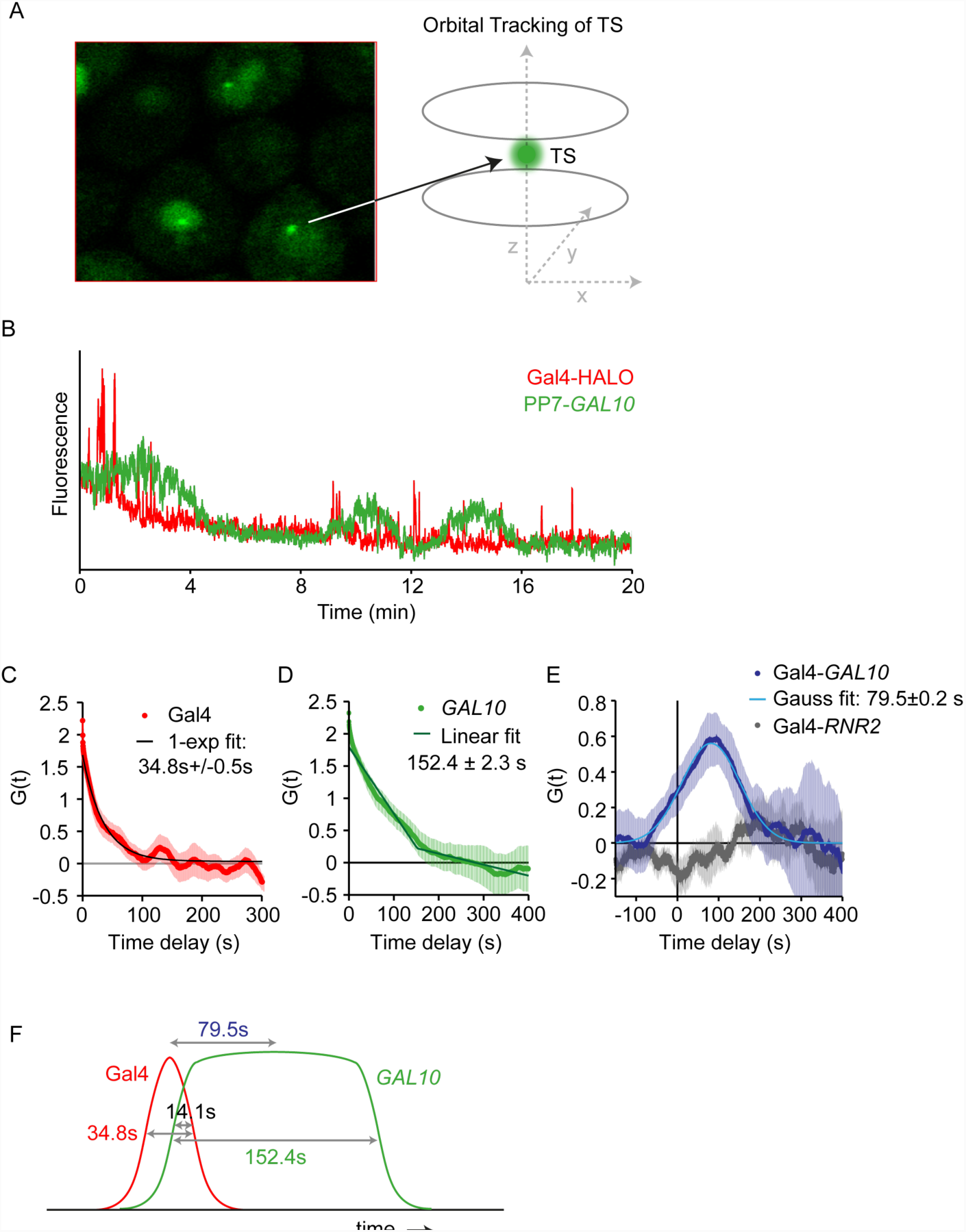
Real-time correlation of Gal4 binding and transcription *in vivo* using orbital tracking. (A) Schematic of 3D orbital tracking of *GAL10* TS. (B) Example trace of Gal4 binding and *GAL10* transcription in the same cell. (C) Autocorrelation of Gal4. The exponential fit shows an average dwell time of 34.8 ± 0.5s. (D) Autocorrelation of *GAL10*. The fit of 2 linear lines reveals a burst duration of 152.4 ± 2.3s. (E) Cross correlations for Gal4-*GAL10* RNA (blue) and Gal4-*RNR2* RNA (grey, negative control). Asymmetry and positive temporal shift of the cross correlation function is consistent with a 79.5 ± 0.2 second delay between the middle of Gal4 and *GAL10* signals, which is not seen in the negative control. (F) Schematic of the different signal durations of Gal4 binding and *GAL10* transcription. Gal4 binding overlaps with the *GAL10* transcription for 14.1s.

The autocorrelation function of Gal4 revealed a dwell time for Gal4 of 34.8s ± 0.5s (Figure 5C), which is longer than the 12s dwell time of the SMT experiments but is likely explained by an underestimation of the dwell time at *GAL10* in the SMT from the stringent colocalization threshold. *GAL10* transcription shows a duration of 152.4s ± 2.3s (Figure 5D), which is comparable to our previously reported dwell time (133s) (Lenstra et al., 2015). The orbital tracking method is thus able to simultaneously capture both Gal4 binding and *GAL10* transcription events.

With the ability to record TF binding and transcriptional activation at the same locus, we are able to measure the temporal sequence of events. The cross-correlation between Gal4 and *GAL10* revealed a positive peak at 79.5s ± 0.2s (Figure 5E), which indicates that fluctuations arising from Gal4 binding and *GAL10* RNA production are temporally correlated and that Gal4 binding precedes the PP7-GAL10 signal by ~80s. Given the dwell time of Gal4 binding, the burst duration of *GAL10* and the 80s delay between the middle of both signals, we calculated that Gal4 binding precedes RNA appearance, but that the nascent RNA levels already start rising during this dwell time with an overlap of 14.1s (Figure 5F). This overlap between Gal4 binding and *GAL10* RNA appearance agrees with our model that several RNA polymerases initiate while Gal4 is bound to the promoter. The temporal correlation was specific, as the cross-correlation peak disappeared in a negative control strain, where a non-galactose responsive gene *RNR2* was tracked together with Gal4 (Figure 5E). In addition, there was no positive correlation in a strain when no synthetic dye was added to the cells to label Gal4 (data not shown). The orbital tracking experiments thus indicate that that the dynamics of Gal4 binding directly determines the *GAL10* transcriptional bursting kinetics.

### Galactose signaling regulates burst frequency, but not burst duration

Gal4 dwell time and the burst duration both depend on the sequence-specific nature of the UAS, which is stably encoded in the genome. However, we asked whether the Gal4 dwell time and the burst duration are also regulated in different conditions, for example during the metabolic response to galactose. Due to the positive and negative feedback loops in the signaling pathway, the galactose responsive genes are known to display a bimodal dose response (Acar et al., 2005; Venturelli et al., 2012), where the fraction of induced cells increases in higher doses of galactose. However, whether the induced cells in different doses of galactose also show different bursting properties is unexplored.

We again focused on *GAL10* transcription to determine burst duration and burst frequency at different galactose levels, because of its higher dynamic range as compared to *GAL3*. Traces of the transcription site intensity showed continuous bursting after the gene was induced (Figure 6A). For lower doses of galactose, the onset of the first burst of *GAL10* transcription was delayed (Figure 6B). Since the off periods were very short and a new burst often started before the previous burst had finished, a threshold to determine the transcription on and off time would result in underestimation of the frequency and overestimation of the burst duration. The autocorrelation of the fluorescence intensity traces was therefore used to extract the burst duration (Figure 6C). In full induction conditions (2% galactose), the burst duration of ~ 95s (Figure 6D) determined from autocorrelation was shorter than the burst duration of ~150s measured previously with the orbital tracking experiments. The reasons for this discrepancy are not known at present but may be due to slightly different bleaching rates between the orbital tracking and time-lapse imaging methods. Nevertheless, across four different galactose concentrations, the burst duration is invariant (Figure 6D), suggesting that the number of initiating polymerases per burst is the same within the resolution of our measurement. Changes in the autocorrelation amplitude also allows for extraction of relative frequencies (Figure 6C). In higher galactose concentration, cells show a higher burst frequency (lower amplitude, Figure 6E), indicating that bursting of *GAL10* is modulated by galactose at the frequency level.

**Figure 6.**
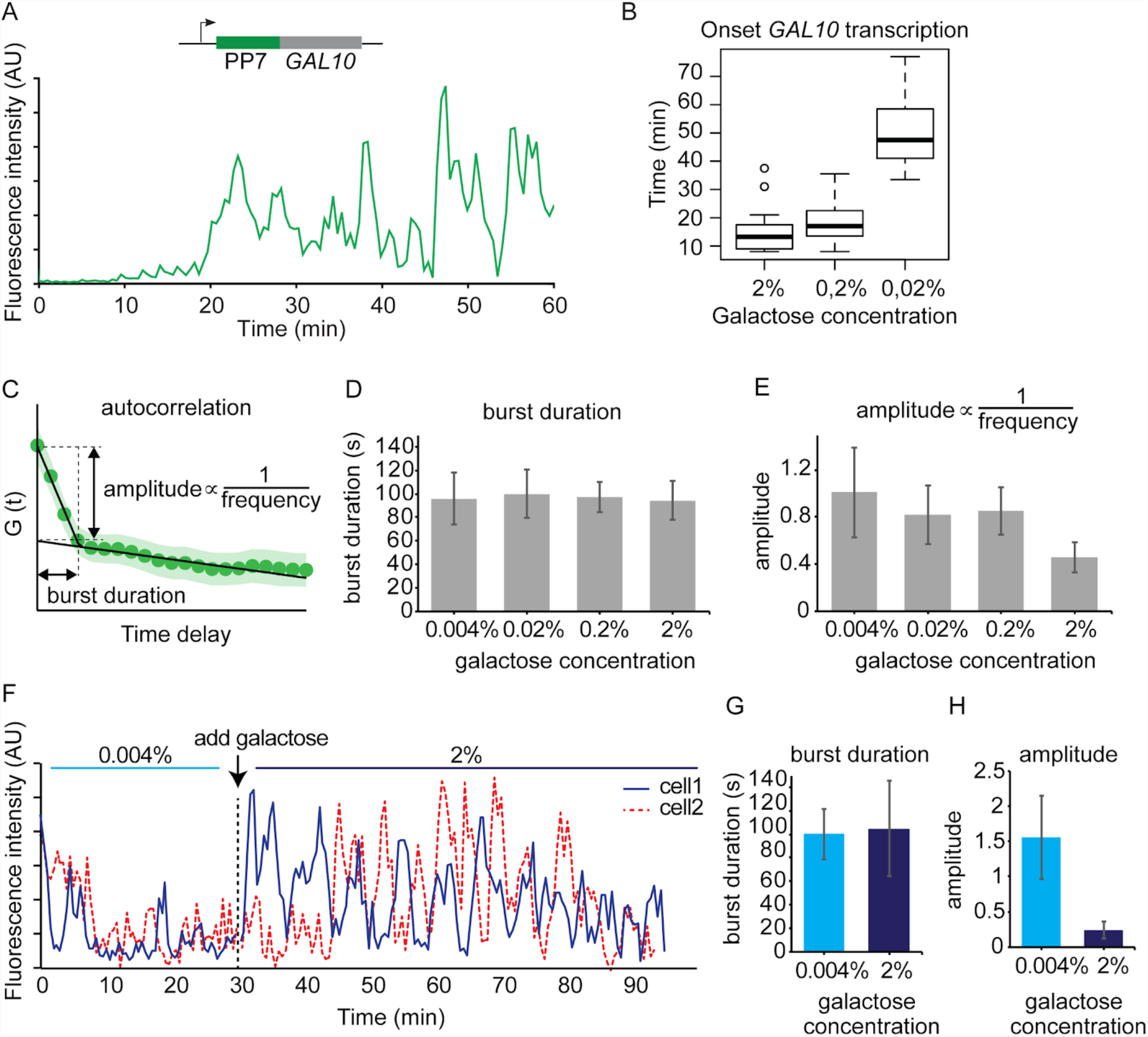
Galactose signaling regulates transcription levels by modulating burst frequency but not burst duration. (A) Example trace of PP7-*GAL10* transcription in 2% galactose. (B) Boxplot of the onset of the first transcription event in different doses of galactose. (C) The autocorrelation was used to interpret the burst duration and burst frequency changes. Burst duration is measured by the intersect of the two linear fits. When burst duration is constant, the amplitude is inversely related to the burst frequency. (D) Burst duration from autocorrelation is constant across 4 different galactose concentrations. (E) Amplitude of autocorrelation decreases (burst frequency increases) at higher doses of galactose. (F) Example traces of two single cells exposed to two different doses of galactose. Same-cell dose response shows same burst duration (G) but lower amplitude (H) at higher galactose concentration, indicating that galactose levels regulate burst frequency.

The burst frequency changes of *GAL10* is about 2-fold. However, the variability between cells is substantial (Figure 6E), making it challenging to compare bursting changes between different conditions. To circumvent this limitation and unambiguously measure transcriptional bursting changes in response to galactose, transcription was monitored in real-time as the galactose concentration is increased during the experiment (Figure 6F). This single-cell dose response is not affected by global cell-to-cell differences, allowing for direct measurements of the bursting changes in the same cell in two different doses of galactose. Again, the burst duration is similar in both concentrations, but the burst frequency clearly increases after increasing the galactose concentration (Figure 6F-H). In summary, we observe no change in burst duration in different galactose concentrations, indicating that the burst duration is not actively regulated by the galactose sensing pathway. Instead, the galactose network regulates transcription levels by modulating the burst frequency and the onset of transcription.

## Discussion

In this study, we have used a combination of *in vivo* and *in vitro* single-molecule imaging approaches to determine the interplay between TF dwell time, nucleosomes and the burst size of a gene. The direct visualization of Gal4 binding and *GAL10* transcription reveals for the first time the timing and correlation of TF binding and target gene expression at single molecule resolution (Figure 5). The observed temporal coupling indicates that the dynamics of TF binding directly determines transcriptional bursting kinetics. Based on our results, we propose a model where multiple RNA polymerases are recruited to form a burst of transcription during the time that TF is bound to the promoter. The burst terminates when the TF dissociates from the DNA (Figure 7).

**Figure 7.**
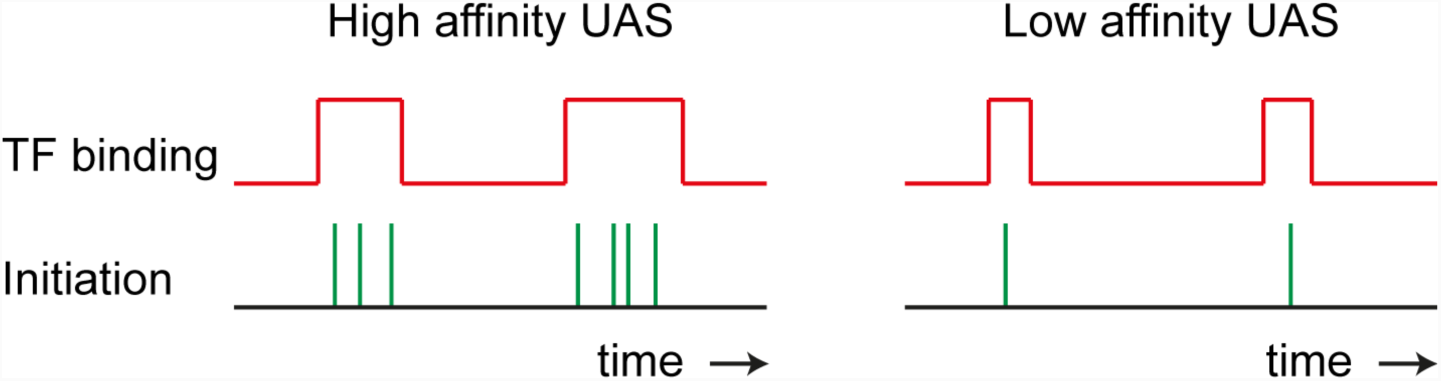
Schematic of transcriptional bursting regulation by TF dynamics. Longer TF binding results in a larger burst size.

Moreover, our data show that the Gal4 dwell time depends on the affinity of the binding site. Mutating the UAS promoter sequence (UASmut) reduces the Gal4 dwell time (Figure 2) and the RNA burst size (Figure 1) but does not affect the Gal4 binding frequency nor the burst frequency. Gal4 dwell time *in vivo* also appears to be considerably reduced by fragile promoter nucleosomes. The *in vivo* average dwell time of 17s (Figure 4) is highly similar to the Gal4 *in vitro* dwell time at nucleosomal DNA (5s, Figure 2), which is remarkable given that many more factors could influence Gal4 binding. In contrast, the Gal4 *in vitro* dwell time at naked DNA is 3 orders or magnitude more stable (Luo et al., 2014), indicating that addition of a nucleosome reduces the dwell time of Gal4 substantially. Indeed, Gal4 is able to bind to nucleosomal templates (Taylor et al., 1991), and at both the *GAL3* and *GAL10* promoter, the Gal4 binding sites appear to be covered by a fragile nucleosome in activating conditions (Figure 3 and (Floer et al., 2010)). Nucleosomes thus not only serve to prevent TF binding, but also appear to play a novel role in limiting the dwell time of TFs. The presence of highly dynamic fragile nucleosomes in many yeast promoters (Kubik et al., 2015) and at many galactose responsive genes (Figure S3) suggests that nucleosomes may be generally required for rapid TF turnover. This mechanism of regulation may only apply to specific TF subclasses that can bind to partially unwrapped nucleosomal DNA, since previous studies in other systems have reported that the position of promoter nucleosomes affect the transcriptional burst frequency (Brown et al., 2013; Dadiani et al., 2013), likely by regulating DNA accessibility and TF binding frequency.

Transcriptional analysis at different galactose doses shows that burst duration is constant in different galactose concentrations (Figure 6). Since the burst duration is coupled to the Gal4 dwell time, the unchanged burst duration along the dose response suggest that Gal4 dwell time is likely not regulated by the galactose signaling network. Instead, galactose regulates the burst frequency. Burst frequency regulation by upstream signaling networks has been observed previously in other systems and is often accompanied by increased transcription factor concentrations (Larson et al., 2013; Senecal et al., 2014). Absolute Gal4 levels do not increase in galactose, but our data is consistent with a model where the signaling network regulates the effective concentration of active Gal4. In inactive conditions Gal4 is still bound to the promoter but repressed by the repressor Gal80 (Sellick et al., 2008), and increasing the galactose concentration could result in dose-dependent unmasking of the Gal4 activation domain by Gal80.

We observe a reduced dwell time of Gal4 in raffinose versus galactose conditions. In raffinose, galactose responsive genes are either lowly transcribed (*GAL80* and *GAL3*) or not transcribed (*GAL10, GAL1* and *GAL7*), and transcription levels are increased in galactose. Even at inactive genes in raffinose, the promoter is bound by Gal4, but inactivated by Gal80. In addition to repressing transcriptional activation by Gal4, the reduced Gal4 dwell time in raffinose (Figure 4E) suggest that Gal80 may possibly act to reduce Gal4 dwell time on DNA. Alternatively, the difference in dwell time of Gal4 could be a result of post-translational modifications (Loffreda et al., 2017), such as Gal4 phosphorylation or ubiquitination (Sellick et al., 2008). It will be interesting to uncover which other factors regulate the dwell time of Gal4.

Live-cell tracking of individual Gal4 molecules by SMT indicates that several second binding of Gal4 is associated with the active transcription site (Figure 4). Dynamic association of Gal4 to chromatin has also been shown by previous competition ChIP experiments (Collins et al., 2009). However, the timescale accessible by ChIP-based approaches is on the order of minutes, whereas SMT experiments report dwell times in the order or seconds. Indeed, the *in vivo* dwell time measurements by single-molecule imaging of TFs is limited by the lifetime of the fluorophore (Liu and Tjian, 2018). If molecules bleach before they dissociate from the DNA, the dwell time can be underestimated or a long-lived binding population can even be entirely missed. Two lines of evidence argue against this scenario for Gal4. First, increasing the frame interval did not increase the dwell time measurements for Gal4 (Figure S4). Second, the positive crosscorrelation between Gal4 and *GAL10* RNA (Figure 5) excludes the possibility that Gal4 would be bound stably to the promoter (on the order of minutes) and be independent from RNA output. Even if the *in vivo* dwell time of Gal4 is underestimated, the peak in the crosscorrelation between Gal4 and *GAL10* would not be consistent with a model where several-minute binding would be required for RNA production.

Recent studies have proposed a model where clustering or phase separation of the transcription machinery are important for driving transcription (Cisse et al., 2013; Hnisz et al., 2017; Lu et al., 2018). Although we do not exclude this possibility, it is important to note that the kinetics of our data can completely be described by the binding kinetics of individual TFs to single target genes, without need to invoke liquid droplets of multi-molecular assemblies of multiple target genes. The correlation between the *in vitro* dwell time and *in vivo* transcriptional burst duration, and the observed similarity between the *in vitro* and *in vivo* dwell time of single Gal4 molecules, suggest that clustering does not contribute greatly to the binding kinetics of Gal4.

The connection between binding site affinity and burst size implies that the burst size of a gene is encoded in the genome. Large transcriptional burst sizes are associated with higher transcriptional noise (Raj et al., 2006; Raser and O’Shea, 2004). As a consequence, TF binding sites in the gene promoter do not only encode information on the level of transcription, but also on the associated cell-to-cell variability. We speculate that the depletion of consensus binding sites and the enrichment of clusters of weaker affinity sites in the genome are perhaps a way to regulate transcription levels without creating large burst sizes. On the other hand, the burst size may also be inherently limited for some genes, as introducing a stronger affinity binding site at *GAL3* did not increase transcription levels (Figure 2K), although the explanation for this phenomenon is still unclear. Perhaps other regulatory factors limit the Gal4 dwell time *in vivo*, or alternatively, the stability of other factors may become limiting when Gal4 dwell time is increased.

In summary, we have used advanced single-molecule microscopy experiments of the galactose system to show how endogenous TFs and nucleosomes regulate bursting kinetics. Whether the mechanism we propose also applies to other transcription networks remains to be determined. Overall, our results provide a framework for interpreting the fluctuations of TF binding and RNA production that have implications for understanding noise in gene expression.

## Acknowledgements

We thank L. Lavis for providing synthetic HALO dyes and G. Mehta, T.S. Karpova, V. Wosika, S. Pelet, J. Wisniewski, Y. Fondufe-Mittendorf, K. Luger, J. Widom, for strains and plasmids. We thank T.S. Karpova for assistance with SMT experiments, Wim de Jonge for help with MNase-seq experiments, Yi Luo for help with the ensemble *in vitro* FRET measurements, and D. Stavreva and members of the D.R.L., M.G.P., M.L.F. and T.L.L. labs for helpful discussions. This work was supported by the Intramural Research Program of the NIH, National Cancer Institute, Center for Cancer Research (D.R.L.), the Research Corporation for Science Advancement and the Gordon and Betty Moore Foundation (GBMF5263.10 M.L.F.), National Institutes of Health (1R15GM123446-01 M.L.F., T32GM086252 B.T.D., R01GM121858 M.G.P, and R01GM121966 M.G.P.), the KWF Dutch Cancer Society (KWF 2012-5394 T.L.L), the Cancer Genomics Center (CGC.nl, T.L.L.) and the European Research Council (ERC Starting Grant 755695 BURSTREG, T.L.L.).

## Author contributions

Conceptualization: T.L.L., M.L.F, D.R.L., M.G.P.; Investigation: B.T.D., A.H. and T.L.L.; Methodology and Software: D.B., M.L.F and T.L.L.; Writing - Original draft: B.T.D., M.L.F. and T.L.L.; Writing - Review and Editing: D.R.L., M.G.P.; Supervision; D.R.L., M.G.P., M.L.F, T.L.L.

## STAR Methods

### KEY RESOURCES TABLE

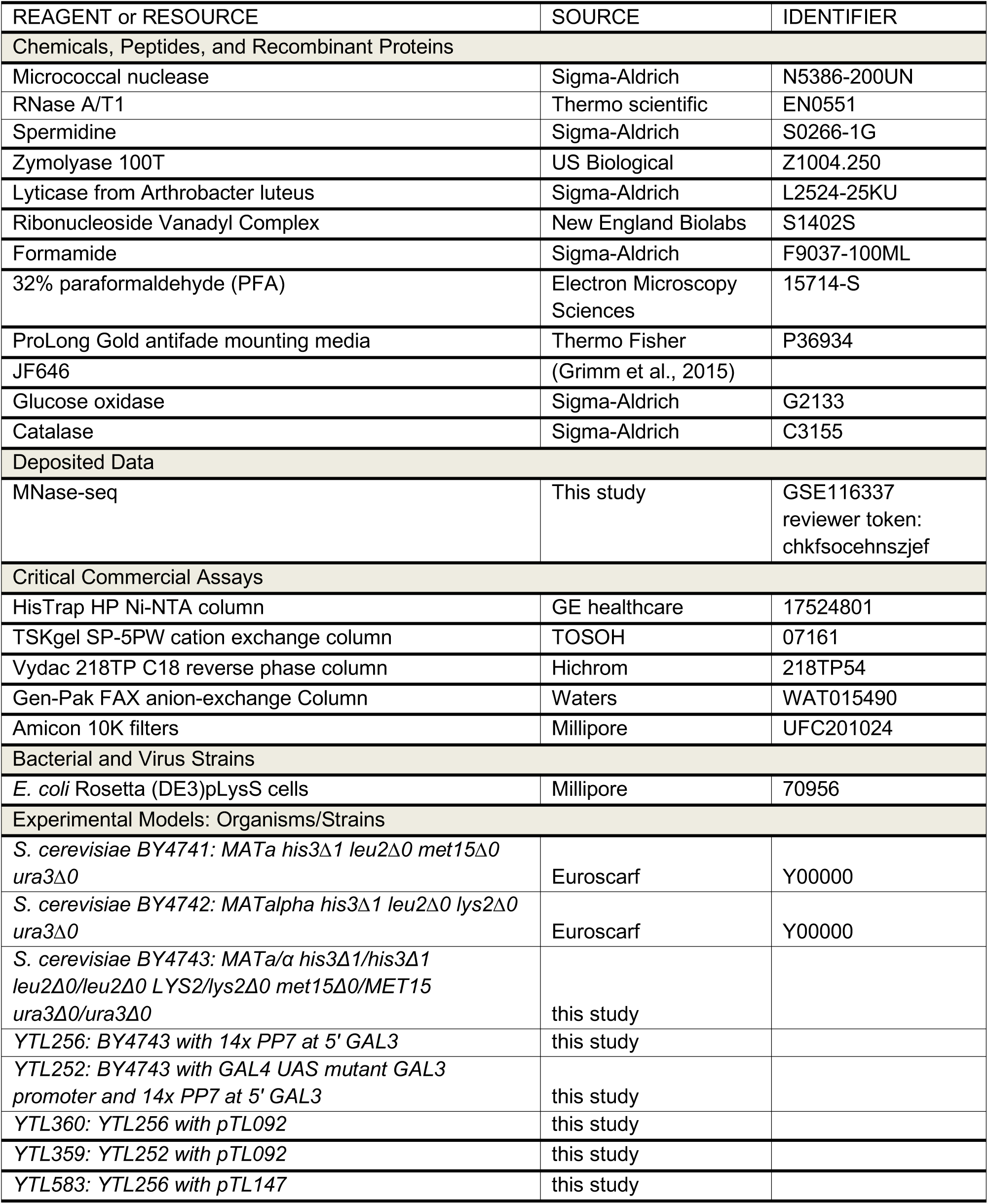
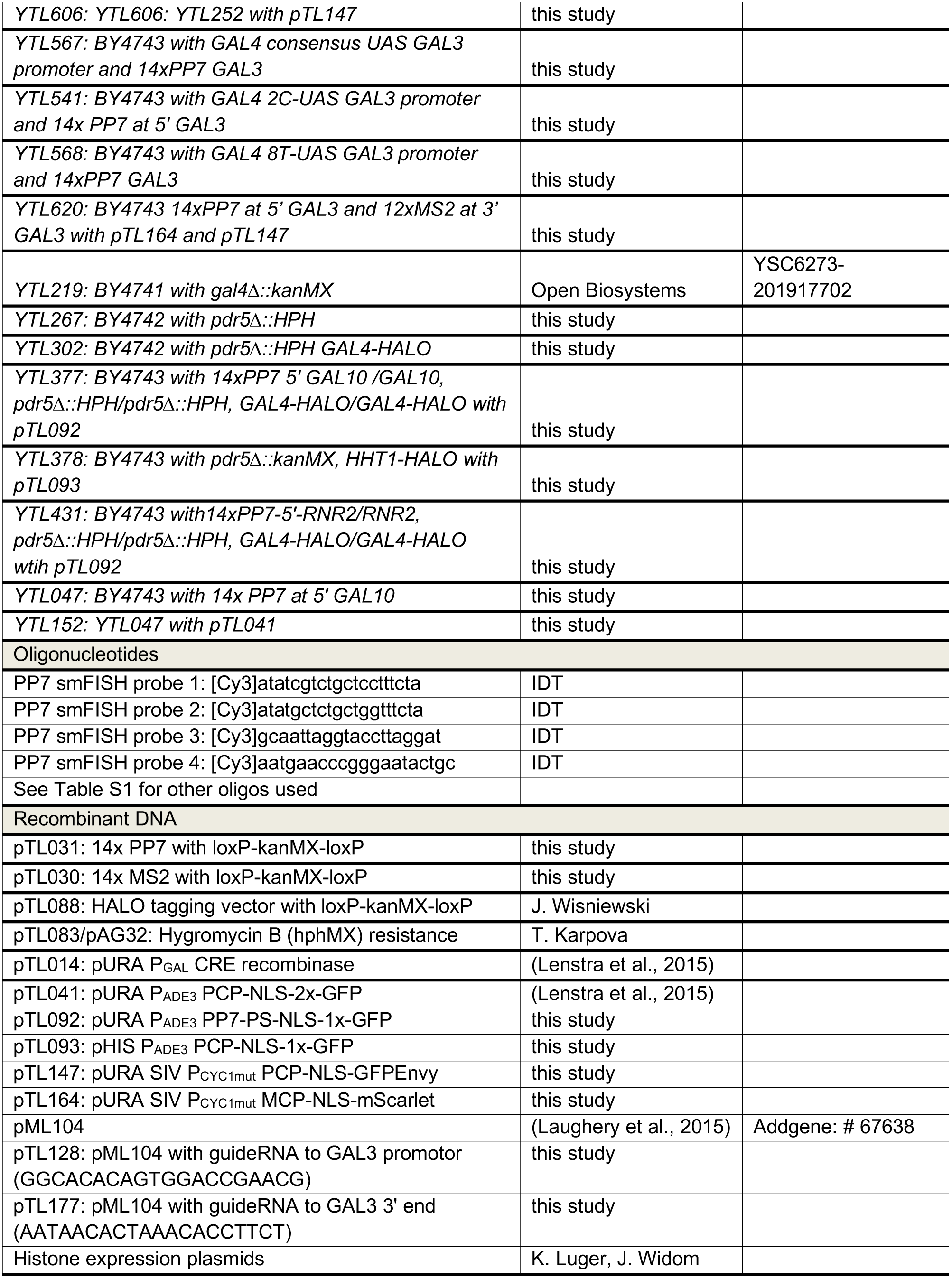
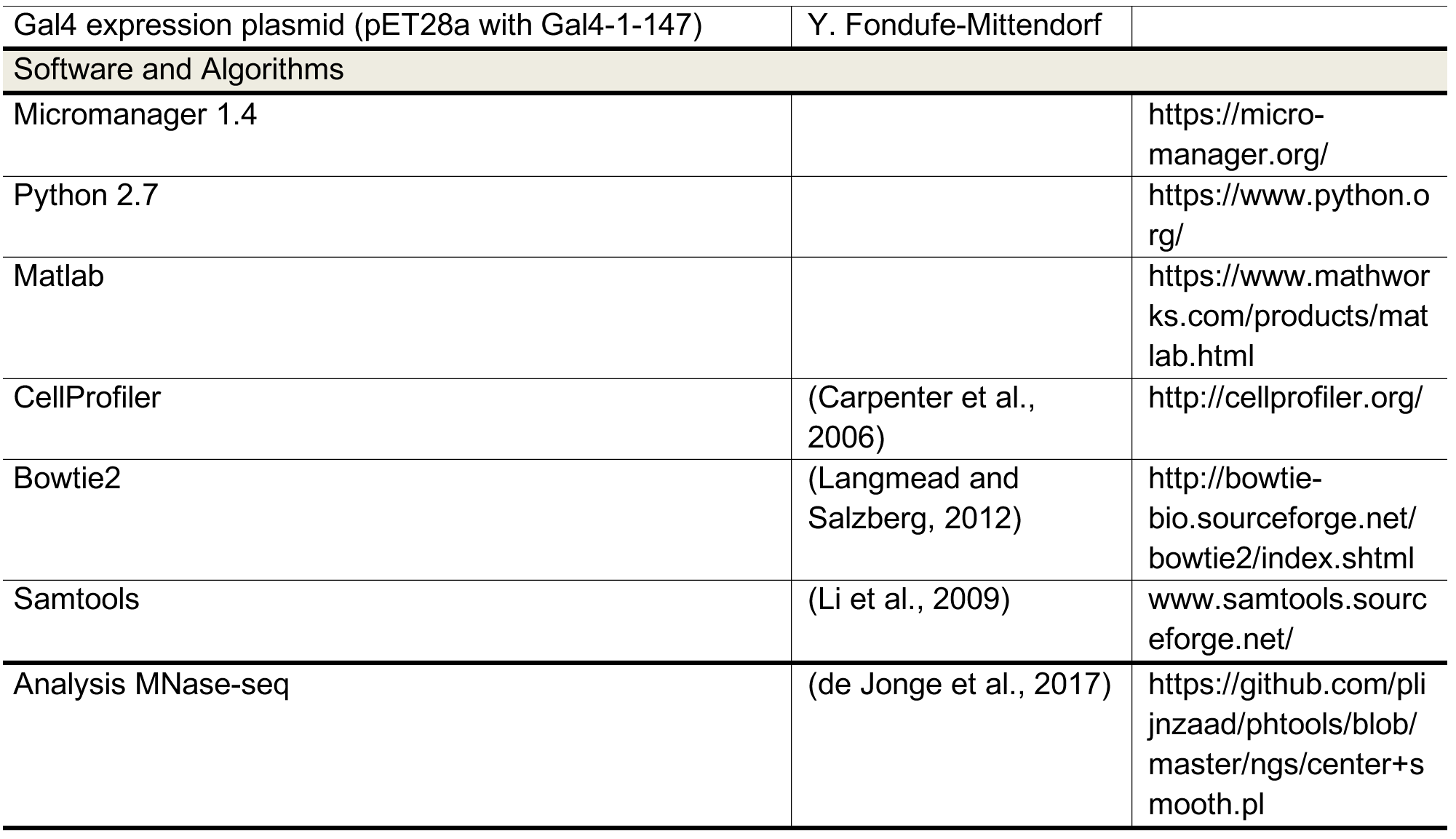

## Method details

### Yeast strains and plasmids

Haploid or diploid yeast was transformed with a PCR product containing the PP7 loop cassette and loxP-kanMX-loxP. The kanMX marker was removed with CRE recombinase (pTL014). Coat protein were expressed from plasmids (pTL041, pTL092, pTL093) or were integrated (pTL147, pTL164). To introduce UAS mutations or the MS2 loop cassette, a PCR product/single stranded oligo and plasmid expressing Cas9 and a guideRNA (Laughery et al., 2015) were transformed. Strains, plasmids and oligos used to construct the strains are listed in the key resource table and Table S1.

### Microscopy of transcription and image processing

Cells were imaged at mid-log (OD 0.2-0.4) on coverslip with 2% agarose pads at 30°C. Imaging of transcription was performed on custom build wide-field microscopes, consisting of an AxioObserver inverted microscope (Zeiss), a 100x NA 1.46 objective, an Evolve 512 EMCCD camera (Photometrics) or a sCMOS ORCA Flash 4v3 (Hamamatsu), a Tokai Hit stage incubator (INUB-LPS) or a UNO Top stage incubator (OKOlab) at 30°C and laser excitation at 488 (Excelsior, Spectra Physics) or LED excitation at 470/24 and 550/15 (SpectraX, Lumencor). Wide-field images of were recorded at 30s interval for PP7-*GAL3* and PP7-*GAL10*, and at 15s interval for PP7-*GAL3*-MS2, with 9 *z*-stacks (δ*z* 0.5 μm) and 150 ms exposure using micromanager software.

For image analysis maximum intensity projections were computed. The intensity of the TS was calculated for each color separately by fitting a 2D Gaussian mask after local background subtraction as described previously (Coulon et al., 2014) using custom Python software. To determine the on and off periods, a threshold was applied to background subtracted traces of 2.5 times the standard deviations of the background. Auto- and crosscorrelation functions were computed and averaged as described previously (Coulon et al., 2014; Lenstra et al., 2015). The burst duration is given by the intersection of the fast and slow linear components.

### Single-molecule FISH

Yeast cultures were grown to early mid-log, fixed with 5% PFA for 20 min, washed 3 times with buffer B (1.2M sorbitol and 100mM potassium phosphate buffer pH 7.5), permeabilized with 300U of lyticase and washed with buffer B. Cells were immobilized on poly-L-lysine coated coverslip and permeabilized with 70% ethanol overnight. Coverslips were hybridized for 4h at 37°C with hybridization buffer containing 10% dextran sulfate, 10% formamide, 2xSSC and 5 pmol probe. For FISH targeting the repeats, four PP7 probes labeled with Cy3 were targeted to the loops (key resource table). For FISH targeting *GAL3*, 48 probes labeled with Quasar 570 were used (Table S1). Coverslips were washed 2x for 30 min with 10% formamide, 2xSSC at 37°C, 1x with 2xSSC, and 1x for 5 min with 1xPBS at room temperature. Coverslips were mounted on microscope slides using ProLong Gold mounting media with DAPI. Cells were imaged on an AxioObserver inverted microscope (Zeiss) with LED illumination (SpectraX, Lumencor) and a sCMOS ORCA Flash 4v3 (Hamamatsu). Spots were localized in maximum intensity projected imaged using custom Python software by fitting to a 2D gaussian mask with local background subtraction. Cell and nuclear outlines were determined with Cell Profiler. TS were defined as the brightest nuclear spot and were normalized to the median fluorescent intensity of cytoplasmic RNAs. The TS distribution was compared with a Poisson distribution. Colocalization of PP7 and *GAL3* probes was determined with a maximum spot distance of 1 pixel.

### Preparation of DNA, nucleosomes and Gal4 for affinity measurements *in vitro*

To prepare DNA molecules for DNA binding experiments, Cy3 or Cy5 oligonucleotides were annealed by mixing at an equal-molar ratio and purified on a gen-pak FAX anion exchange column (Waters). To prepare DNA molecules for nucleosome experiments, oligonucleotides for PCR were labeled with Cy3 NHS ester (GE healthcare) at an amino group at their 5’-ends. Oligonucleotides were then purified by HPLC with 218TP C18 reverse phase column (Hichrom). DNA molecules were prepared by PCR from a plasmid containing the 601 nucleosome positioning sequence (NPS) with a UASwt or UASmut Gal4 binding site positioned 8 bp into the nucleosome. Following PCR amplification, DNA molecules were purified using a MonoQ column (GE healthcare). All DNA sequences can be found in Table S1.

Human recombinant histones were expressed and purified as previously described (Luger et al., 1999). Mutation H3(C110A) was introduced by site-directed mutagenesis (Agilent). The histone octamer was refolded by adding each of the histones together at a ratio of 1.1:1.1:1:1 (H2A:H2B:H3:H4) and purifying as previously described (Luger et al., 1999). H2A(K119C) containing HO was labeled with Cy5-maleamide (GE Healthcare) as previously described (Gibson et al., 2016).

Nucleosomes containing Cy3-labeled DNA and purified Cy5-labeled histone octamer were reconstituted through double dialysis. DNA and octamer were mixed at a ratio of 1.25:1 DNA:HO in 0.5x TE pH 8 with 1 mM benzamidine hydrochloride (BZA) and 2M NaCl in a volume of 50 uL. The mixture was loaded into a dialysis chamber and placed into a large dialysis bag containing 80 mL of 0.5x TE pH 8, 1 mM BZA, and 2M NaCl and dialyzed extensively against 0.5x TE pH 8 with 1 mM BZA at 4C. Dialyzed nucleosomes were loaded onto 5-30% sucrose gradients and purified by centrifugation on an Optima L-90 K Ultracentrifuge (Beckman Coulter) with a SW-41 rotor. Sucrose fractions containing nucleosomes were collected, concentrated, and stored in .5x TE pH 8 on ice.

To prepare Gal4, amino acids 1-147 of Gal4 was expressed from plasmid pET28a containing Gal4-1-147 in *E. coli* Rosetta (DE3)pLysS cells by inducing with 1 mM IPTG + 10 uM ZnAc for 3 hours. Cells were harvested by centrifugation and resuspended at 50 mL per 1 L starting culture in buffer A (50 mM Tris pH 8.0, 200 mM NaCl, 1 mM DTT, 10 uM ZnAc, 1 mM phenylmethanesulfonyl fluoride (PMSF), 20 ug/mL leupeptin, and 20 ug/mL pepstatin) and stored at -80 C. The cells were lysed by sonication and clarified by centrifugation, loaded onto a 5 mL HisTrap HP Ni-NTA column, equilibrated with buffer B (25 mM Tris pH 7.5, 200 mM NaCl, 0.2% Tween-20, 10 mM imidazole, 20 uM ZnAc, 1 mM DTT, 1 mM PMSF) and eluted with elution buffer (25 mM Tris pH 7.5, 200 mM NaCl, 0.2% Tween-20, 200 mM imidazole, 20 uM ZnAc, 1 mM DTT, 1 mM PMSF). Fractions containing Gal4 were dialyzed into buffer C (25 mM Tris pH 7.5 200 mM NaCl, 20 uM ZnAc, 1 mM DTT, 1 mM PMSF) and purified with a TSKgel SP-5PW cation exchange column (Tosoh). Fractions containing Gal4 were concentrated using Amicon 10K filters (Millipore) and stored in buffer D (HEPES pH 7.5, 200 mM NaCl, 10% glycerol, 20 uM ZnAc, 1 mM DTT, 1 mM PMSF).

### Ensemble PIFE measurements

Gal4 binding to its target site on a 51 bp Cy3/5-DNA was determined by protein induced fluorescence enhancement (PIFE) (Hwang et al., 2011), where Cy3 fluorescence increases upon protein binding. Fluorescence spectra were acquired with a Fluoromax4 fluorometer (Horiba) using an excitation wavelength of 510 nm. Gal4 affinity to DNA was measured by incubating 0.5 nM DNA with 0-30 nM Gal4 in 10 mM Tris-HCl pH 8, 130 mM NaCl, 10% glycerol, 0.0075% v/v Tween-20. From these experiments, we determined that ~80% of DNA molecules are bound by ~5 nM Gal4. Competition experiments were performed by incubating Cy3/5 DNA with 5 nM Gal4 and 0-500 nM unlabeled competitor DNA containing either the UASwt or UASmut Gal4 binding site for 30 minutes. For each titration we determined the S1/2, the concentration at which the fluorescence signal decreases by 50%. With these measurements, we calculated the relative affinity of Gal4 to the UASwt and UASmut sequences (relative affinity 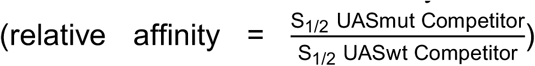. Fluorescence spectra were analyzed with Matlab to determine the change in Cy3 fluorescence.

### Ensemble FRET measurements

Gal4 binding to its site within a nucleosome was detected through FRET. 0.5 nM nucleosomes were incubated for at least 5 minutes with 0-1000 nM Gal4 in 10 mM Tris-HCl pH 8, 130 mM NaCl, 10% glycerol, 0.0075% v/v Tween-20. FRET efficiency was calculated using the RatioA method with a custom matlab program (Clegg, 1992).

### Single molecule TIRF microscope

The smTIRF microscope was built on an inverted IX73-inverted microscope (Olympus) as previously described (Roy et al., 2008). 532 and 638 nm diode lasers (Crystal Lasers) were used for Cy3 and Cy5 excitation. The excitation beams were expanded and then focused through a quartz prism (Melles Griot) at the surface of the quartz flow cell. A 1.3 N.A. silicone immersion objective (Olympus) was used to collect fluorescence which were separately imaged onto an iXon3 EMCCD camera (Andor) with a custom built emission path containing bandpass filters and dichroic beam splitters (Chroma Tech). Each video was acquired using Micro-Manager software.

### Single molecule fluorescence measurements of Gal4 binding kinetics

Flow cells were functionalized as previously described (Kinz-Thompson et al., 2013). Briefly, quartz microscope slides (Alfa Aesar) were sonicated in toluene and then ethanol, and then further cleaned in piranha solution (3:1 mixture of concentrated sulfuric acid to 50% hydrogen peroxide). Slides were washed in water and, once completely dry, incubated in 100 uM mPEG-Si and biotin-PEG-Si (Laysan Bio) overnight in anhydrous toluene. Functionalized quartz slides and coverslips were assembled into microscope flow cells using parafilm to define channels. Before each experiment, the flow cell is treated sequentially with 1 mg/ml BSA, 40 ug/ml streptavidin, and biotin-labeled nucleosomes.

Biotinylated nucleosomes were allowed to incubate in the flow cell at room temperature for 5 minutes and then washed out with imaging buffer containing the desired concentration of Gal4. The samples were first exposed to 638 nm excitation to determine the location of Cy5 molecules and then 532 nm for FRET measurements. The imaging buffer for FRET experiments contained 10 mM Tris-HCl pH 8, 130 mM NaCl, 10% glycerol, 0.0075% v/v Tween-20, 0.1 mg/ml BSA, 2 mM Trolox, 0.0115% v/v COT, 0.012% v/v NBA, 450 ug/ml glucose oxidase and 22 ug/ml catalase.

Single molecule time series were fit to a two-state step function by the hidden Markov method using vbFRET (Bronson et al., 2009). Idealized time series were further analyzed using custom written Matlab programs to determine the dwell-time distributions of the TF bound and unbound states. Approximately 50% of FRETing traces fluctuated and were used in the analysis of FRET data. Dwell-time and unbound-time cumulative sum distributions were fit to single-exponential distributions using matlab to obtain rate constants for the transitions between bound and unbound states.

### MNase-seq

Preparation and analysis of mono-nucleosomal DNA was performed as described previously (de Jonge et al., 2017) with minor modifications. Briefly, cells were grown in SC + 2% raffinose or SC + 2% galactose from OD 0.3 to OD 1.0, fixed in 1% PFA, washed with 1M sorbitol, treated with spheroplasting buffer (1M sortibol, 1mM β-mercaptoethanol, 10 mg/ml Zymolyase 100T) and washed twice with 1M sorbitol. Spheroplasted cells were treated with 0.01171875 or 0.1875 U micrococcal nuclease in digestion buffer (1M sorbitol, 50 mM NaCl, 10 mM Tris pH7.4, 5 mM MgCl_2_, 0.075% NP-40, 1 mM β-mercaptoethanol, 0.5 mM spermidine) at 37°C. After 45 min, reactions were terminated on ice with 25 mM EDTA and 0.5 % SDS. Samples were treated with proteinase K for 1h at 37°C and decrosslinked overnight at 65°C. Digested DNA was extracted with phenol/chloroform (PCI 15:14:1), precipitated with NH_4_-Ac, and treated with 0.1 mg/ml RNaseA/T1. The extent of digestion was check on a 3% agarose gel.

Sequencing was performed on one HiSeq2500 lane using Illumina TruSeq v4 chemistry. Paired-end 125 bp reads were aligned with Bowtie 2. Nucleosome dyads were found by taking the middle of each paired read of insert size between 95 and 225 bp, and were smoothed with a 31 bp window, as described in (de Jonge et al., 2017).

### Growth assay

Serial dilutions (5-fold) of BY4741, YTL219, YTL267 and YTL302 strains were spotted on YEP + 2% glucose, YEP + 2% galactose + 20 ug/ul Etidium Bromide, and YEP + 2% raffinose + 2% galactose + 40 mM lithium chloride + 0.003% methionine. Growth was assessed after 3 days at 30°C.

### Single molecule tracking of Gal4

Cells were grown in SC + 2% glucose overnight, washed in SC + 2% raffinose, and grown in SC+ 2% raffinose for 4h. After 2h, cells were labeled with 5 nM JF646 (for Gal4) and nM (for H3). Before imaging, cells were washed once and immobilized on coverslip with 2% agarose pads containing 2% raffinose or 2% galactose. SMT movies were acquired a custom-built microscope, containing a Highly Inclined Laminated Optical (HILO) sheet illumination at 488nm and 647 nm, a 100x 1.49 TIRF objective (Olympus), three EM-CCD iXon Ultra 888 cameras (Andor), and an UNO stage top incubator at 30°C (OKOlab). Images captured simultaneously with dual-color illumination at 488 and 647 nm for 30 ms at 200 ms or 1000 ms interval.

Spots were tracked independently for the *GAL10* transcription site and Gal4 channel using custom Matlab software based on MatTrack (Mazza et al., 2013). Particle positions were fit with a 2D gaussian mask, tracked using nearest-neighbors approach, and manually verified. Only molecules that were tracked in more than 4 frames were considered bound. To determine whether a molecule was bound or diffusing, a threshold was used based on tracking of histone H3. Tracking of histone H3 (*HHT1*-HALO) showed that 99% of single molecules had a frame-to-frame displacement of less than 0.35 μm at 200 ms interval, or less than 0.37 μm at 1000 ms interval. These maximum displacements were used to determine if Gal4 particles are chromatin bound or diffusing. The cumulative distribution of dwell times of bound Gal4 molecules (survival probability plot) was corrected for bleaching by dividing the distribution by the decay of the number of particles found over time. To extract the average dwell times, the survival probability distribution was fit to a one or two-component exponential decay function.

To select for traces that colocalized with the *GAL10* transcription site, images were registered using beads, and a 250 nm distance threshold was used to determine overlap. The survival probability plot was fit to a one-component dwell exponential function.

### Orbital tracking

Cells were prepared as for SMT experiments. Active TSs were identified at high laser power, but once orbital tracking of an TS was initiated, the power was reduced to reduce photobleaching. Orbital tracking was performed according to (Kis-Petikova and Gratton, 2004; Levi et al., 2005a, 2005b). Tracking of transcription sites was achieved using two orbits of radius ~87nm at a position 145nm above the TS, followed by two orbits 145nm below the TS, with 64 points per orbit and a pixel dwell time of 1024 µs per pixel. Each orbit had a duration of 65.5ms with a total sampling time of 262 ms (3.8 Hz) sampling rate, allowing for the measurement of Gal4 and *GAL10* RNA occupancy at high sampling rate for 10-20 minutes while keeping the transcription site in focus via the z-piezo and active feedback from microscope software.

Imaging and orbital tracking were performed on an ISS Alba FCS microscope (Champaign) with 488nm and 633nm excitation, two SPCM-ARQH Avalanche Photodiodes (Pacer), a 1MHz IOtech 3000 Data Acquisition card (Measurement Computing Corporation,) and a Nano-F25HS high speed z piezo (Mad City Labs) coupled to a Nikon Ti-U inverted microscope with a CFI Plan Apochromat 60X 1.2 NA water immersion objective (Nikon). Data acquisition was performed using SimFCS 3.0 software written by Enrico Gratton (Laboratory for Fluorescence Dynamics, University of California, Irvine). Bead alignment showed that the point spread function alignment in both channels was ~60nm. When using one photon excitation for cross correlation spectroscopy, alignment of both lasers in 3D is critical to achieve success (Schwille et al., 1999). A 0.5-2x variable beam expander was therefore placed in front of the 488nm laser to achieve alignment in z due to chromatic aberration.

Fluorescence intensity appeared in carpet plots of angle vs. time showing a Gaussian peak in the RNA channel with intermittent signal in the red channel assumed to be binding of fluorescently labelled Gal4 molecules. Sections of the fluorescence intensity traces were selected for active transcription by the appearance of signal above background in the carpet plots. Data analysis was performed using custom software written in IDL (Harris Geospatial Solutions) and Python. Temporal correlation functions of the fluorescent signal were calculated as previously from 1/2 the average photon intensity over the 4-orbital period of 262ms directly from the DC component of the fluorescence intensity trace as shown in equation 18 of (Kis-Petikova and Gratton, 2004). Correlation functions were averaged over 10-20 measurements. This resulted in very robust and reproducible correlation functions from which the dwell time of Gal4 molecules, *GAL10* RNA and temporal relationships could be estimated. The *GAL10* RNA autocorrelation is best fit with a linear model as reported in (Larson et al., 2011), Gal4 autocorrelation function is best fit using an exponential reaction dominant diffusion binding model (Digman and Gratton, 2009; Michelman-Ribeiro et al., 2009) and the cross correlation is best fit to a shifted Gaussian model.

